# Investigating discriminating concentrations for monitoring susceptibility to broflanilide and cross resistance to other insecticide classes in *Anopheles gambiae* sensu lato, using the new WHO bottle bioassay method

**DOI:** 10.1101/2022.10.04.510874

**Authors:** Renaud Govoetchan, Abibath Odjo, Damien Todjinou, Graham Small, Augustin Fongnikin, Corine Ngufor

## Abstract

**Background:** Broflanilide is a new insecticide being developed for malaria vector control. As new insecticide chemistries become available, strategies to preserve the susceptibility of local malaria vectors and extend their useful life need to be considered before large scale deployment. This requires the development of appropriate testing procedures and identification of suitable discriminating concentrations for monitoring susceptibility in wild vector populations to facilitate decision making by control programmes.

**Methods:** Dose-response WHO bottle bioassays were conducted using the insecticide-susceptible *Anopheles gambiae* s.s. Kisumu strain to determine a discriminating concentration of broflanilide. Bioassays were performed without the adjuvant Mero^®^ and with two concentrations of Mero^®^ (500 ppm and 800 ppm) to investigate its impact on the discriminating concentration of the insecticide. Probit analysis was used to determine the lethal doses at 50% (LC_50_) and 99% (LC_99_) at 24-, 48- and 72-hours post-exposure. Cross-resistance to broflanilide and pyrethroids, DDT, dieldrin and carbamates, was investigated using *An. gambiae* s.l. Covè and *An. coluzzii* Akron strains. The susceptibility of wild pyrethroid-resistant mosquitoes from communities in Southern Benin to broflanilide was assessed using the estimated discriminating concentrations.

**Results:** Broflanilide induced a dose-dependent and delayed mortality effect. Mortality rates in bottles treated without Mero^®^ were <80% using the range of broflanilide doses tested (0-100 µg/bottle) leading to high and unreliable estimates of LC_99_ values. The discriminating concentrations defined as 2XLC_99_ at 72 hours post exposure were estimated to be 2.2 µg/bottle with 800 ppm of Mero^®^ and 6.0 µg/bottle with 500 ppm of Mero^®^. Very low resistance ratios (0.6-1.2) were determined with the insecticide resistant *An. gambiae* s.l. Covè and *An. coluzzii* Akron strains suggesting the absence of cross-resistance via the mechanisms of resistance to pyrethroids, DDT, dieldrin and carbamates they possess. Bottle bioassays performed with broflanilide at both discriminating concentrations of 6 µg/bottle with 500 ppm of Mero^®^ and 2.2 µg/bottle with 800 ppm of Mero^®^, showed susceptibility of wild highly pyrethroid-resistant *An. gambiae* s.l. from villages in Southern Benin.

**Conclusion:** Here we determined discriminating concentrations for monitoring susceptibility to broflanilide in bottle bioassays, using susceptible *An. gambiae* vectors. Using the estimated discriminating concentrations, we showed that wild pyrethroid-resistant populations of *An. gambiae* s.l. from southern Benin were fully susceptible to the insecticide. Broflanilide also shows potential to be highly effective against *An. gambiae* s.l. vector populations that have developed resistance to other public health insecticides.

## 1. Introduction

Since 2015, the rate of progress in reducing malaria cases and deaths has stalled in many countries with moderate or high transmission [1]. Several factors have contributed to this slowdown in progress, including the development of vector resistance to major public health insecticides. Insecticides from five classes of compounds (pyrethroids, organochlorines, carbamates, organophosphates and neonicotinoids) have been recommended for malaria vector control [2]. Resistance to pyrethroids is now widespread which seriously limits the effectiveness of this previously ideal insecticide class for malaria vector control [3, 4]. Some vector populations have developed resistance to multiple insecticides [3, 5–7]; a disturbing situation necessitating the development of new effective insecticide chemistries with different modes of action that can provide improved control of insecticide resistant vector populations and facilitate insecticide resistance management [4, 8].

The meta-diamide, broflanilide, a novel active ingredient with a unique chemical structure developed by Mitsui Chemicals Agro Inc.; Tokyo, Japan [9], is being developed for public health use. This insecticide acts as a non-competitive antagonist (NCA) of the γ-aminobutyric acid (GABA) receptor of chloride channels of the insect inhibitory nervous system [10]. Broflanilide (trade name TENEBENAL^™^) is being developed for use on insecticide treated nets (ITNs) and for indoor residual spraying (IRS) against malaria vectors [11, 12]. A wettable powder formulation of this insecticide (VECTRON^TM^ T500) has shown potential to control pyrethroid resistant malaria vectors when applied for indoor residual spraying (IRS) [12, 13]. A recent experimental hut trial in Southern Benin also demonstrated prolonged control of wild pyrethroid resistant *Anopheles gambiae* s.l. with VECTRON^TM^ T500 over 18 months (Govoetchan and Ngufor, unpublished data). Community trials are ongoing in Benin and Tanzania to assess the impact of IRS with VECTRON^™^ T500 when applied on a large scale in an area where vectors are resistant to pyrethroids. Data from these studies are being reviewed by the World Health Organisation’s Vector Control Product Prequalification (WHO PQT/VCP) to evaluate the suitability of VECTRON^™^ T500 for large scale use in IRS.

As new insecticide chemistries are developed for vector control, strategies to preserve the susceptibility of local malaria vectors to these insecticides and extend their useful life need to be considered before large scale deployment [8, 14]. This includes the development of suitable testing procedures for monitoring susceptibility in wild vector populations to facilitate decision making by control programmes [14]. Surveillance for potential resistance to an insecticide in a given vector species requires the identification of a suitable discriminating concentration for testing the susceptibility of wild vector populations to the insecticide in laboratory assays. The discriminating concentration is typically defined as a dose of an insecticide that will reliably kill vector mosquitoes that are susceptible to the insecticide within a defined observation time post exposure. Laboratory bioassays using insecticide discriminating concentrations have historically provided an efficient method for monitoring insecticide resistance in local malaria vector populations to help guide context-specific decision making around their deployment for vector control [15].

The WHO has recently recommended new bottle bioassay procedures and discriminating concentrations for investigating resistance in mosquito vectors to new insecticides that are not suitable for impregnation onto filter papers [14, 16, 17]. The bottle bioassay is a practical method that offers the opportunity to include an adjuvant to help improve the distribution and presentation of insecticides that tend to crystallise on surfaces. Crystallisation can prevent optimal pickup and uptake of the insecticide by exposed mosquito vectors leading to inconsistent results [14, 18]. The addition of an adjuvant may facilitate the identification of a suitable discriminating concentrations for broflanilide and thus needs to be explored. Considering the limited number of insecticide chemistries available for malaria vector control, it is also helpful to investigate the potential for cross-resistance to broflanilide via existing insecticide resistance mechanisms in vector populations as this could limit its usefulness for malaria vector control.

In this study, we performed a series of laboratory bottle bioassays to: *i)* determine a discriminating concentration for monitoring susceptibility to broflanilide, with and without an adjuvant, in *An. gambiae* s.l. mosquitoes using a susceptible strain; *ii)* investigate cross-resistance to broflanilide and other vector control insecticides in this vector species and; *iii)* assess susceptibility of wild *An. gambiae* s.l. in villages in Southern Benin to the estimated discriminating concentrations.

## 2. Methods

### 2.1. Mosquito strains

The following mosquito strains were used for the testing:

- *An. gambiae* sensu stricto Kisumu, an insecticide-susceptible reference strain originating from the Kisumu area in Kenya and was colonized at CREC/LSHTM insectary.
- *An. coluzzii* Akron, a pyrethroid and carbamate resistant strain originating from Akron (9°19′N2°18′E), Southern Benin, and maintained at CREC/LSHTM insectary. Resistance is mediated by target site mutations (L1014F kdr and Ace-1R) and overexpressed cytochrome P450 enzymes [19].
- *An. gambiae* sensu lato (s.l.) Covè, a pyrethroid, DDT and dieldrin resistant strain originating from Covè (7°14′N2° 18′E), Southern Benin. The strain is composed of a mixture of *An. coluzzii* and *An. gambiae* s.s.. Resistance is mediated by a target site kdr mutation (L1014F) and overexpressed cytochrome P450 enzymes [20].

### 2.2. Susceptibility of test strains to insecticides

As part of a quality management system at the CREC/LSHTM insectary, all three mosquito strains (*An. gambiae* s.s. Kisumu; *An. gambiae* s.l. Covè and *An. coluzzii* Akron) were routinely tested every three months to validate their susceptibility status to public health insecticides. Four rounds of routine bioassays were performed during the study. Standard WHO susceptibility cylinder bioassays were performed with 8 different insecticides which included 3 pyrethroids (0.05% deltamethrin, 0.75% permethrin, 0.05% alphacypermethrin), 2 organochlorines (4% DDT, 4% dieldrin), 2 organophosphates (0.1% fenitrothion, 0.25% pirimiphos-methyl) and 1 carbamate (0.1% bendiocarb) [21]. All treated papers were obtained from Universiti Sains Malaysia, Malaysia. Tests were conducted with unfed 2-5 days old mosquitoes of each strain. They were exposed for 1h to insecticide-treated papers and mortality was recorded 24h later. Approximately 100 mosquitoes (four replicates of 25 mosquitoes) were used per insecticide and mortality was pooled across the different time points. Control mosquitoes were exposed to untreated papers. Bioassays were performed at a temperature of 27 °C ± 2 °C and a relative humidity of 75% ± 10%.

### 2.3. Dose response bottle bioassays with broflanilide

To determine the discriminating dose of broflanilide, we performed dose response bottle bioassays with the susceptible *An. gambiae* ss Kisumu strain[21]. The bottles were treated with eight (8) doses of broflanilide ranging between 0 and 10 µg/bottle (0, 0.05, 0.1, 0.25, 0.5, 1, 2.2, 4.6, and 10 μg/bottle). Given broflanilide’s tendency to crystalise, the bottle treatments were done using a mixture of acetone and the adjuvant Mero^®^ (81% rapeseed oil methyl ester) to improve insecticide pickup and uptake. To help identify a suitable dose for Mero^®^ for the bottle bioassays, for each broflanilide dose, bottles were treated using two different concentrations of Mero^®^ (500 ppm and 800 ppm). Bottles treated with each insecticide dose and acetone alone (without addition of Mero^®^) were also prepared. Four replicate bottles were prepared per treatment following WHO procedures [16]. One hundred (100) unfed adult female *An. gambiae* Kisumu (2-5 days old) were exposed for 1 hour to each treatment in cohorts of 25 mosquitoes per bottle. After exposure, the mosquitoes were gently aspirated from the bottle into clean, labelled paper cups and provided with 10% sugar solution soaked in cotton wool. Knock down was recorded after exposure, and mortality was recorded at 24 hours intervals up to 72 hours post-exposure. Mosquitoes were also exposed to bottles treated with acetone and Mero^®^ alone applied at 500ppm and 800ppm to serve as controls. The bioassays were conducted at 27 °C ± 2 °C and a RH of 75% ± 10%. Lethal doses inducing 50% (LC_50_) and 99% (LC_99_) mortality at 24h, 48h and 72 hours post-exposure were determined for each concentration of Mero^®^ using probit analysis. The discriminating concentration of broflanilide with each dose of Mero^®^ was determined by doubling the LC_99_ in line with WHO procedures [14].

### 1.4 Investigating cross-resistance to insecticides used in vector control

To investigate the potential for cross-resistance to mechanisms providing resistance to pyrethroids, DDT, dieldrin and carbamates, dose-response bottle bioassays were conducted with the *An. gambiae* s.l. Covè and *An. coluzzii Akron* strains and compared to the susceptible *An. gambiae* s.s. Kisumu strain. For these bioassays, nine (9) doses of broflanilide ranging between 0 and 10 μg/bottle (0, 0.05, 0.1, 0.25, 0.5, 1, 2.2, 4.6, and 10 μg /bottle) were tested with addition of Mero^®^ at 800 ppm. The LC_50_ were determined for each strain using probit analysis. Resistance ratios relative to the susceptible Kisumu strain were also determined.

### 2.5. Susceptibility of wild pyrethroid-resistant *An. gambiae* s.l. from Southern Benin to broflanilide

To investigate the susceptibility of wild pyrethroid resistant malaria vectors from communities in Southern Benin to broflanilide at the discriminating concentrations determined in this study, we performed bottle bioassays with adult female mosquitoes (2-5 days) emerging from larvae collected in 5 villages of the sub-district of Za-Kpota Centre, Southern Benin: Za-Kékéré, Dokpa, Za-Zounmè, Kèmondji and Détèkpa. For each village, 100 F1 mosquitoes were exposed for 1 hour and knockdown was recorded after exposure and mortality every 24 hours up to 72 hours post-exposure. Bottle bioassays were also conducted with deltamethrin at 1X, 2X, 5X and 10X the discriminating concentration (12.5 μg/bottle) to determine the intensity of pyrethroid resistant in wild *An. gambiae* sl vector populations from each village.

### 2.6. Data analysis

Data from bottle bioassays were subjected to log-probit regression analysis using SPSS statistics 28.0v and LC_50_ and LC_99_ were calculated with 95% confidence intervals. Susceptibility of broflanilide in the wild vector populations was determined from the percentage mortality, following World Health Organization (WHO) guidelines on insecticides susceptibility: mortality ≥ 98% indicates susceptibility, mortality less than 90% indicates the existence of resistance while if the observed mortality is between 90 and 97%, the presence of resistant genes in the vector population must be confirmed [21].

## 3. Results

### 3.1. Susceptibility of mosquito strains

The pooled mortality rates of insectary-maintained *An. gambiae* s.s. Kisumu, *An. coluzzii Akron* and *An. gambiae* s.l. Covè strains exposed to discriminating concentrations of insecticides in WHO susceptibility bioassays are presented in Figure 1. In total 10,617 female mosquitoes were tested in routine WHO cylinder bioassays between February 2020 and December 2021. *An. gambiae* s.s. Kisumu remained susceptible to all insecticides throughout this period. *An. gambiae* s.l. Covè showed resistance to pyrethroids (16-35%), DDT (2%) and dieldrin (80%) and susceptibility to organophosphates and carbamates (99-100%), whilst the *An. coluzzii* Akron strain was resistant to pyrethroids, DDT and bendiocarb and showed suspected resistance to organophosphates (90-97%).

**Figure 1:**
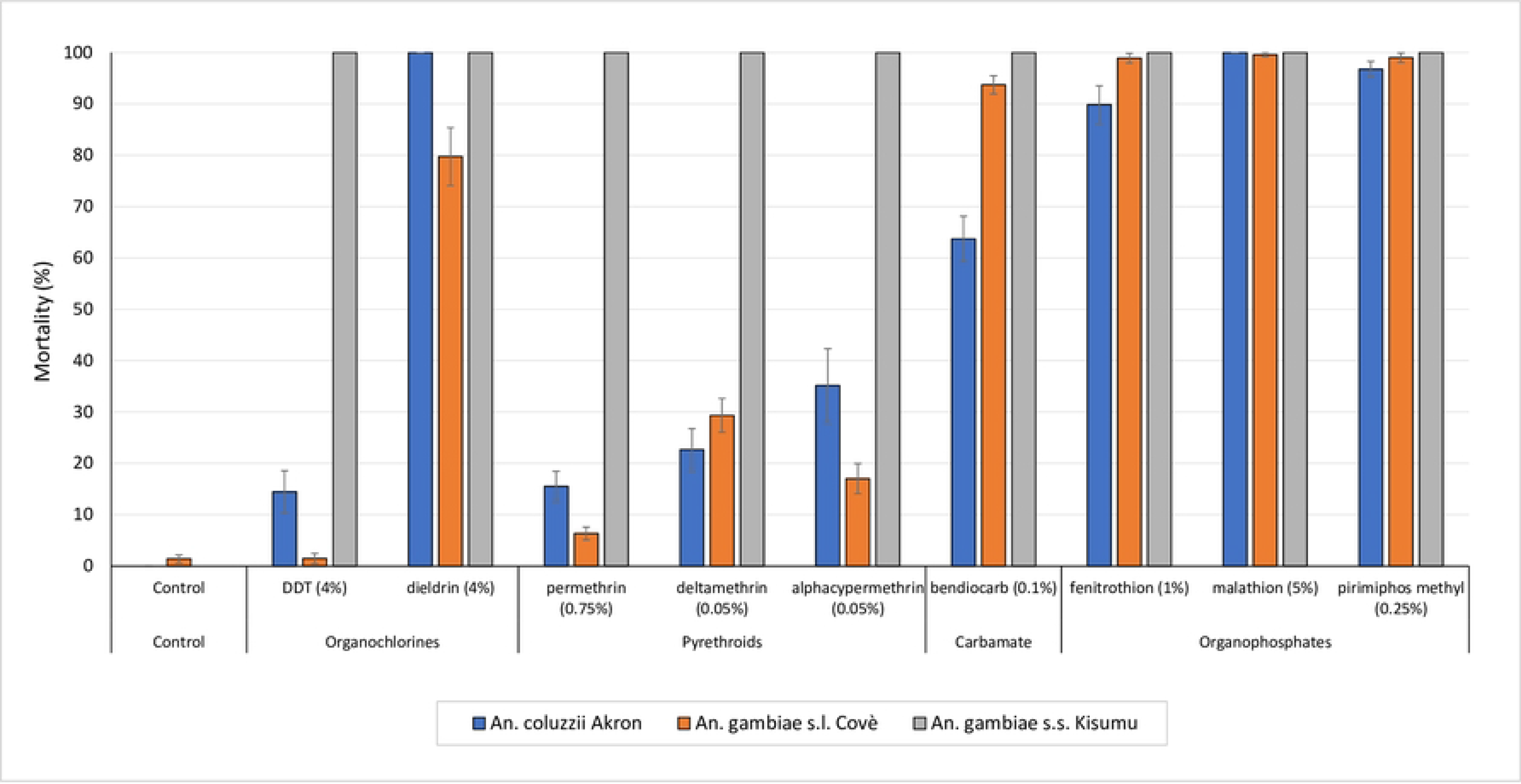
Susceptibility of *An. gambiae* s.s. Kisumu, *An. gambiae* s.l. Covè and *An. coluzzii* Akron exposed to discriminating concentrations of insecticides in WHO cylinder bioassays. Data was pooled from routine testing on insectary-maintained mosquito strains performed between February 2020 and December 2021. Error bars represent 95 % confidence intervals.

### 3.2. Results from dose-response bottle bioassay with insecticide-susceptible *An. gambiae* Kisumu

The results from the broflanilide dose response bottle bioassays performed with the insecticide susceptible *An. gambiae* s.s. Kisumu strain with bottles treated without Mero^®^ and with Mero^®^ (applied at 500 ppm and 800 ppm) are presented in Figure 2 for mortality rates observed and Table 1 for the LC_50_ and LC_99_ and estimated discriminating concentrations. A total of 2,842 female mosquitoes were tested in the dose response bottle bioassays with the susceptible Kisumu strain. Mortality in the untreated control bottles was <5% irrespective of the dose of Mero^®^ added. Knockdown rates were generally low across all treatments tested but this increased as the insecticide dose increased (Supplementary Table S1).

**Figure 2:**
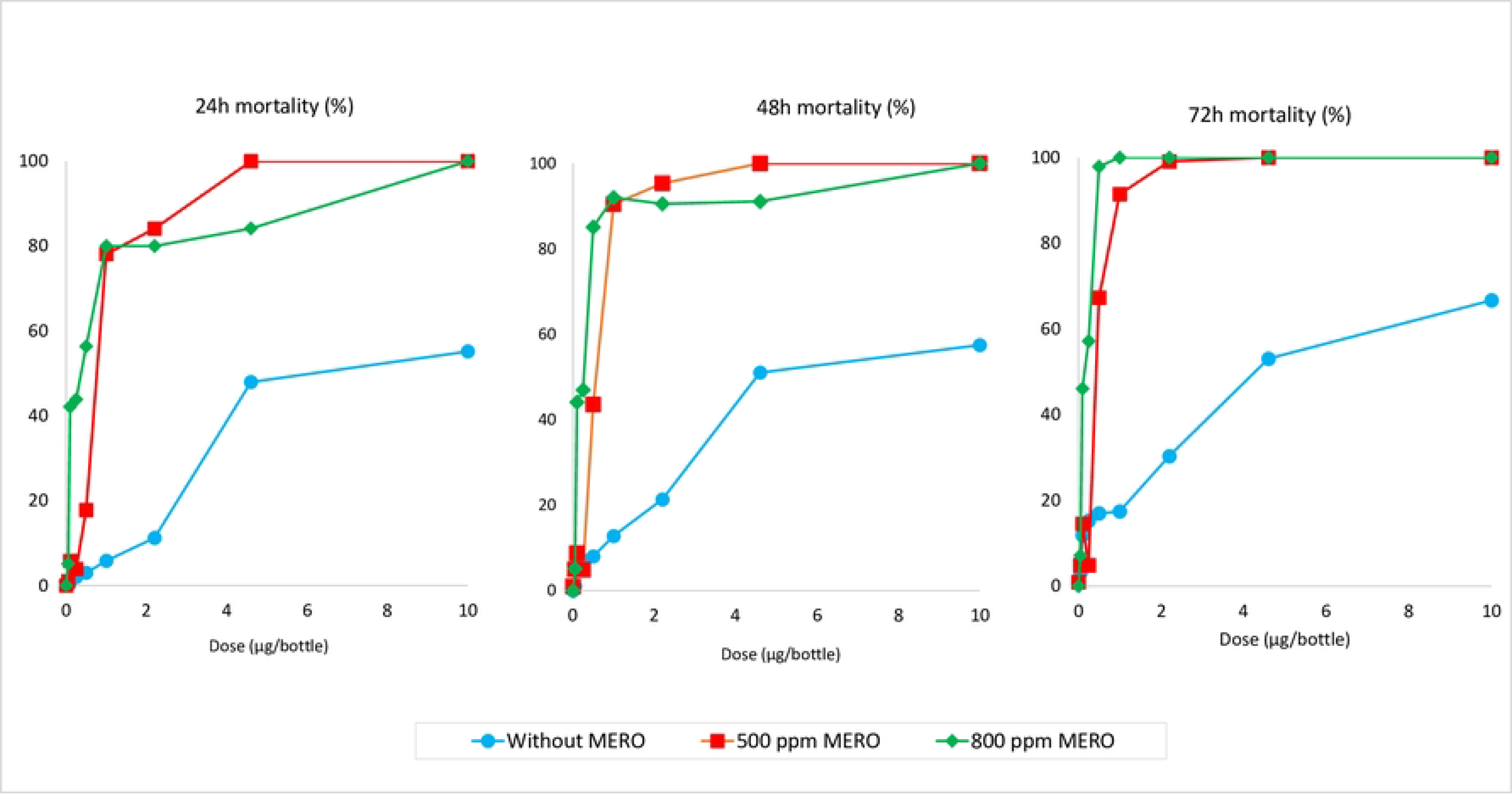
Dose response mortality results with susceptible *An. gambiae* s.s. Kisumu at a) 24h b) 48h and c) 72h post exposure to broflanilide treated bottles with varying doses of Mero^®^ adjuvant.

**Table 1:** Lethal concentrations (LC_50_ and LC_99_) and discriminating concentrations of broflanilide against insecticide susceptible *An. gambiae* s.s. Kisumu in bottle bioassays with and without Mero^®^ adjuvant. Probit analysis was performed using SPSS software.

| Dose of Mero® | Observation time | slope | LC<sub>50</sub><br>(95% CI) | LC<sub>99</sub><br>(95% CI) | Discriminating<br>concentration*<br>(µg/bottle) |
| --- | --- | --- | --- | --- | --- |
| No Mero® | 24 hours | 1.584 | 7.7<br>(6.1 - 10.4) | 246.2<br>(123 - 661) | 492.5<br>(245 – 1322) |
|  | 48 hours | 1.548 | 7.6<br>(5.4 - 11.8) | 1607<br>(558 - 7145) | 3214<br>(1115 – 14288) |
|  | 72 hours | 1.450 | 5.1<br>(3.3 - 9.1) | 2464<br>(581.5 - 2361.3) | 4928<br>(1163 – 4723) |
| Mero® 500 ppm | 24 hours | 5.073 | 0.7<br>(0.6 - 1.0) | 6.0<br>(3.5 - 14.5) | 11.9<br>(7.066 – 29.092) |
|  | 48 hours | 4.935 | 0.5<br>(0.4 - 0.6) | 4.1<br>(2.6 – 8.2) | 8.2<br>(5.2 – 16.3) |
|  | 72 hours | 4.762 | 0.4<br>(0.3 - 0.5) | 3.0<br>(2.0 - 5.5) | 6.0<br>(4.080 – 10.932) |
| Mero® 800 ppm | 24 hours | 2.803 | 0.3<br>(0.1 – 0.7) | 35.5<br>(8.3 – 1210.3) | 71.0<br>(16.6 – 2420.7) |
|  | 48 hours | 2.569 | 0.2<br>(0.1 - 0.4) | 10.0<br>(3.0 -194.0) | 20.1<br>(6.0 – 387.9) |
|  | 72 hours | 2.400 | 0.1wel<br>(0.094 - 0.225) | 1.1<br>(0.5 - 8.1) | 2.2<br>(1.1 – 16.1) |

\*Discriminating concentrations are determined by doubling the LC<sub>99</sub> values

There was a clear relationship between broflanilide dose and mortality in treated bottles (Figure 2). Mortality rates also increased as observation time increased from 24h to 72h showing evidence of a delayed mortality effect. In the absence of Mero^®^, none of the doses tested induced 100% mortality (Figure 2); further tests were performed without Mero^®^ at doses of 21.5μg, 49μg and 100μg but none of these induced >90% mortality. Hence it was not possible to determine a reliable LC_99_ value for broflanilide in bottles treated without Mero^®^ with the range of doses tested. The efficacy of broflanilide improved substantially with addition of Mero^®^ with mortality being higher at a given broflanilide dose in bottles treated with 800 ppm of Mero^®^ than in bottles treated with 500 ppm (Figure 2). The LC_50_ and LC_99_ values were thus lower with bottles treated with Mero^®^ compared to those treated without Mero^®^ and increased as the dose of Mero^®^ increased (Table 1). The discriminating concentrations for broflanilide determined from the LC_99_ values showed a corresponding decrease when bottles were treated with Mero^®^, being 492.5 μg at 24h and 4.9 g at 72h without Mero^®^, 11.9 μg at 24h and 6.1 μg at 72h with Mero^®^ 500 ppm, and 71.0 μg at 24 and 2.2 μg at 72h with Mero^®^ at 800 ppm (Table 1).

### 3.3. Investigation of cross-resistance to pyrethroids, carbamates and organophosphates

Results from the dose response bioassays, using Mero^®^ at 800 ppm, performed with insecticide resistant *An. gambiae* s.l. Covè and *An. coluzzii* Akron strains in comparison to the susceptible *An. gambiae* s.s. Kisumu strain are summarised in Figure 3 for 72h mortality rates and Table 2 for the lethal concentrations and resistance ratios obtained. A clear dose response relationship with mortality was observed with all three strains (Figure 2). The LC_50_ values were higher with the *An. gambiae* s.l. Covè strain compared to the susceptible *An. gambiae* s.s. Kisumu strain but lower with the *An. coluzzii* Akron strain. The resistance ratios observed were very low (0.64-1.2 fold) with both insecticide resistant strains which suggests the absence of cross-resistance to broflanilide via the insecticide resistance mechanisms present in the Covè and Akron strains.

**Figure 3:**
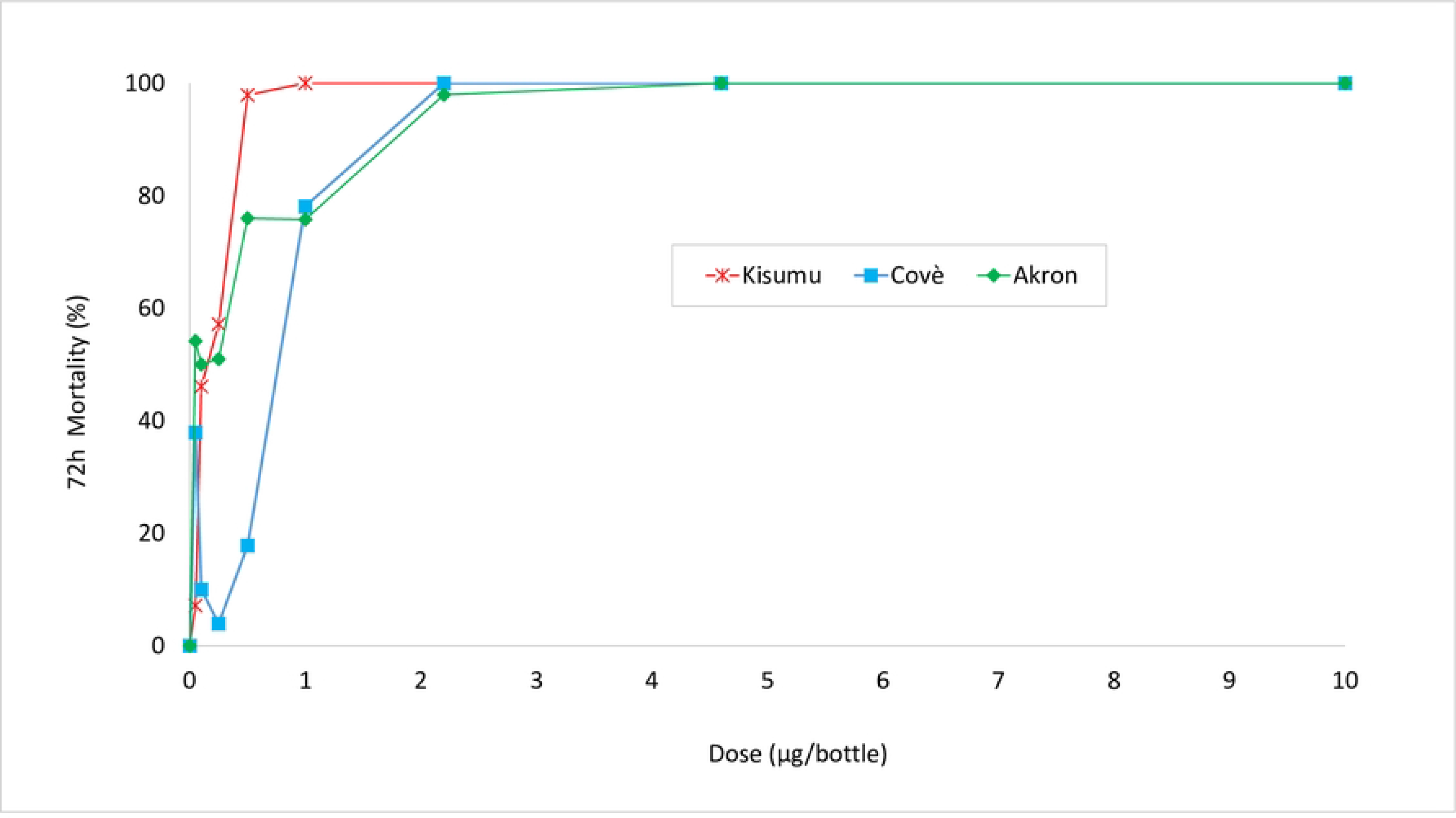
Mortality rates (72h) in broflanilide dose response bottle bioassays with 800 ppm of Mero^®^ with *An. gambiae* s.s. Kisumu, *An. gambiae* s.l. Covè, and *An. coluzzii* Akron strains.

**Table 2:** Lethal concentrations (LC_50_) and resistance ratios of *An. gambiae* s.l. Covè, *An. coluzzii* Akron compared to *An. gambiae* s.s. Kisumu in bottle bioassays with broflanilide. *Probit analysis was performed with 72h mortality data on mosquitoes exposed to bottles treated with broflanilide and Mero*^®^ applied at *800 ppm using SPSS software*.

| Strain | LC<sub>50</sub> | 95% Confidence Intervals | RR<sub>50</sub> |
| --- | --- | --- | --- |
| *An. gambiae* s.s. Kisumu | 0.146 | 0.094 - 0.225 | - |
| *An. gambiae* s.l. Covè | 0.177 | 0.078 - 0.306 | 1.21 |
| *An. coluzzii* Akron | 0.094 | 0.025 - 0.187 | 0.64 |

### 3.4. Results from bottle bioassays with wild pyrethroid-resistant *An. gambiae* s.l. from southern Benin using discriminating concentrations of broflanilide

The deltamethrin resistance intensity bottle bioassays showed a high intensity of resistance to pyrethroids in the selected villages in Southern Benin; mortality rates were <60% at 10 times the discriminating concentration (Figure 4). Bottle bioassays performed with broflanilide at the discriminating concentrations of 6 μg/bottle (with 500 ppm Mero^®^) and 2.2 μg/bottle (with 800 ppm Mero^®^), showed susceptibility to the insecticide in mosquitoes sampled from all 5 villages (Figure 5).

**Figure 4:**
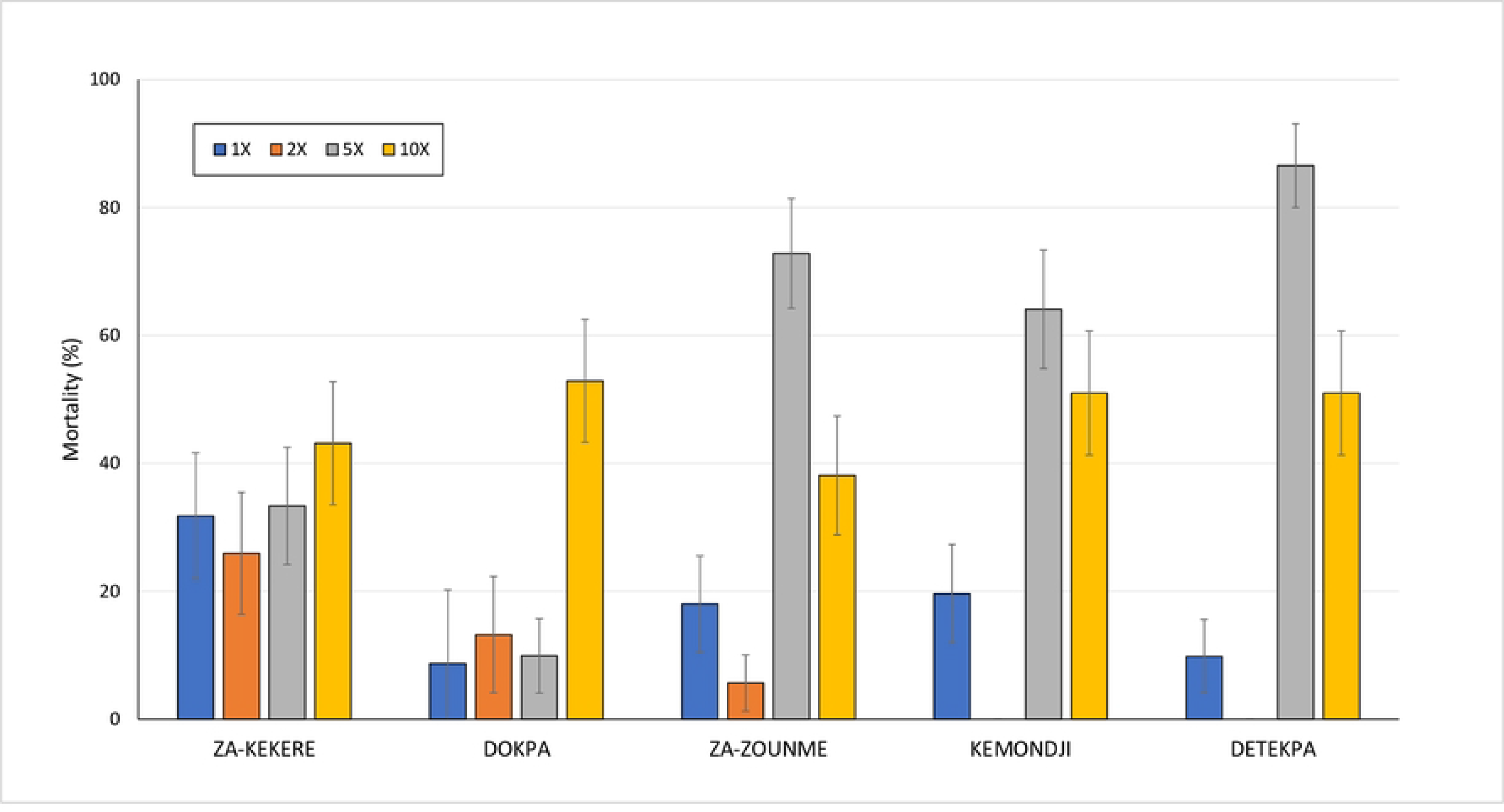
Mortality (24 hours) deltamethrin 1X, 2X, 5X and 10X in bottle bioassays with wild *An. gambiae* s.l. from 5 villages of Za-Kpota sub-district, southern Benin.

**Figure 5:**
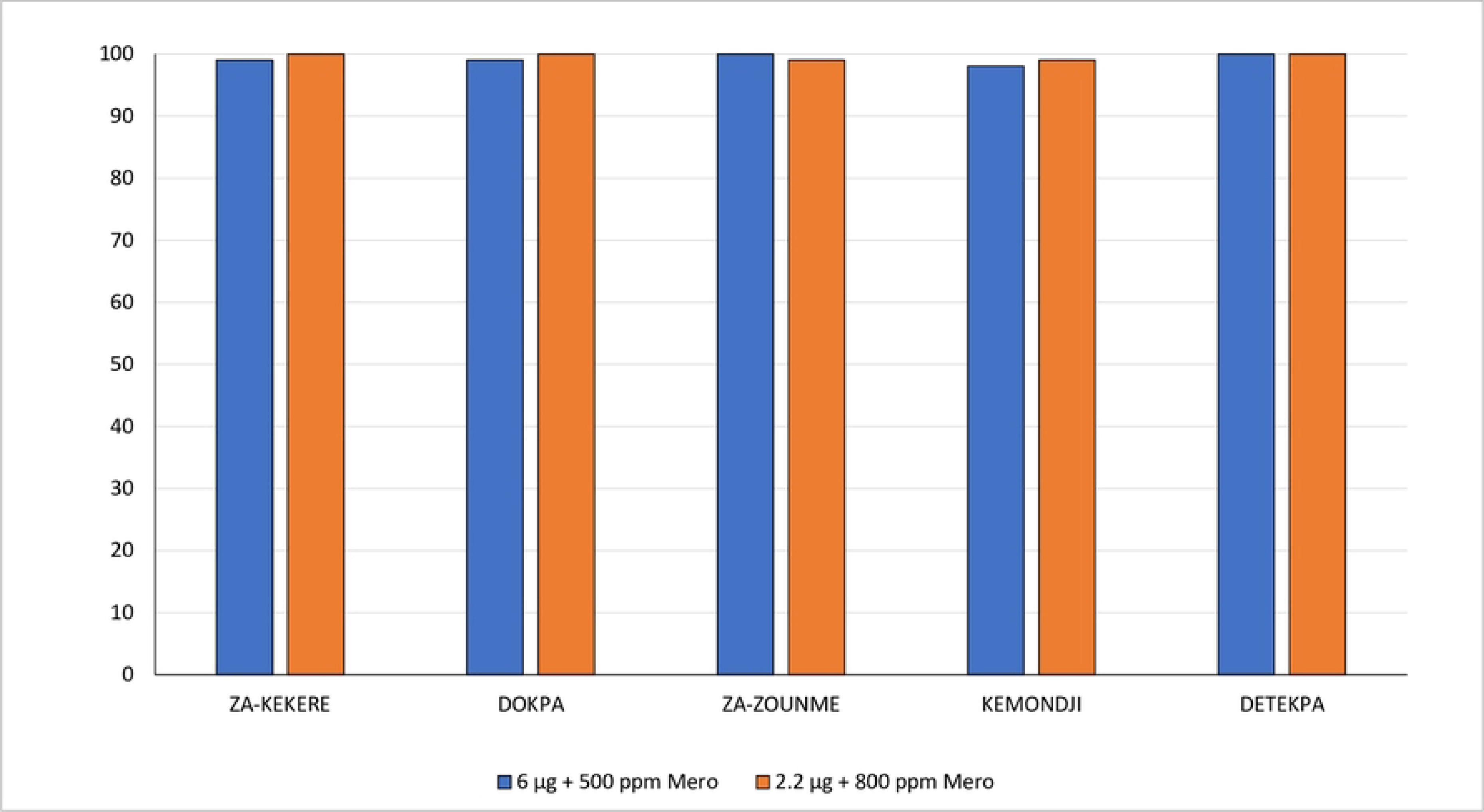
Mortality (72h) of wild *An. gambiae* s.l. from southern Benin in bottle bioassays with estimated discriminating concentrations of broflanilide.

## 4. Discussion

The purpose of this study was to determine a discriminating concentration and identify suitable methods for monitoring susceptibility to broflanilide, a newly discovered meta-diamide insecticide, in wild malaria vector populations using bottle bioassays. We also investigated the potential for cross-resistance to broflanilide through mechanisms of resistance to other insecticides in local *An. gambiae* s.l. vectors. Following previous studies with malaria vectors, we defined the discriminating concentration as double the concentration that was calculated to induce 99% mortality in a susceptible strain [14]. The results showed that determination of a reliable discriminating concentration required the addition of an adjuvant. In the absence of the adjuvant, 100% mortality could not be achieved with any of the doses tested resulting in unusually high and unreliable discriminating concentrations for broflanilide ranging from 429.3 μg at 24hrs post exposure to 4928 μg at 72hrs post exposure. These results suggest that bottle bioassays performed with broflanilide against wild vector populations without an adjuvant might result in misleading reports of resistance to the insecticide. This corroborates previous studies investigating resistance to clothianidin in bottle bioassays without Mero^®^ that showed low potency of the insecticide with widely inconsistent results [18, 22, 23]. In our study, when Mero^®^ was added at concentrations of 500 ppm and 800 ppm, the potency of broflanilide increased, resulting in lower discriminating concentrations at the different observation times post-exposure (24h - 72h). Based on the delayed activity of broflanilide on mosquito vectors requiring an observation time of up to 72hrs post exposure [12], 6 μg with 500 ppm of Mero^®^ and 2.2 μg with 800 ppm of Mero^®^ can be considered suitable discriminating concentrations for broflanilide.

The addition of an adjuvant in bottle bioassays is recommended for insecticides that tend to crystallise on bottle surfaces preventing adequate uptake by exposed mosquitoes [16, 18]. However, the role of adjuvants and their impact on insecticide resistance bottle bioassay endpoints have been controversial. It has been suggested that adding adjuvants in routine bottle bioassays may result in overestimation of vector susceptibility to insecticides due to their underlying toxicity [23]. Previous studies investigating resistance to clothianidin in malaria vectors using higher concentrations of Mero^®^ (1500 ppm) have often reported high levels of control mortality >20% suggesting that the results obtained might be confounded by the toxicity of the adjuvant (unpublished data). Here we observed higher mosquito mortality rates with the higher dose of Mero^®^ compared to the lower dose which demonstrates a dose response effect of Mero^®^ on the bottle bioassay outcome. Nevertheless, mosquito mortality in control bottles treated with acetone and Mero^®^ applied at 500 ppm and 800 ppm were very low (<5%) suggesting these lower doses of MERO^®^ are less likely to confound the bottle bioassay results with broflanilide in routine monitoring bioassays.

A recent WHO-coordinated multi-centre study investigating the discriminating concentration of new public health insecticides, recommended a standard dose of 800 ppm of Mero^®^ for determining resistance in all anopheline vectors (except *An. albimanus*), to insecticides that require the addition of an adjuvant in bottle bioassays [14]. The methods used in our study also followed the new WHO bottle bioassay procedures in terms of bottle coating, drying time, exposure time, endpoints etc [16, 17]. This suggests that the lower discriminating concentration of 2.2 μg applied with Mero^®^ at 800 ppm, as recommended by WHO, could be a more suitable discriminating concentration for monitoring resistance to broflanilide in *An. gambiae* s.l. vectors using bottle bioassays. Nevertheless, we observed full susceptibility of wild highly pyrethroid resistant *An. gambiae* s.l. mosquitoes from 5 selected villages in Benin using both discriminating concentrations of 2.2 μg (with 800 ppm of Mero^®^) and 6.0μg (with 500 ppm of Mero^®^). Based on this finding, where high control mortality rates are observed with Mero^®^ at 800 ppm, it might be useful to consider running broflanilide bottle bioassays with a lower Mero^®^ dose of 500 ppm and the higher discriminating concentration of 6 μg .

Knowledge of insecticide cross-resistance patterns is essential for selecting appropriate insecticides for local contexts and for designing insecticide resistance management strategies. Broflanilide effectiveness would be compromised if cross-resistance were to be detected via existing mechanisms that confer resistance in malaria vectors to other public health insecticides. Both *An. gambiae* s.l. strains tested in the cross resistance bioassays exhibited high levels of resistance to pyrethroids mediated by high frequencies of the L1014F knockdown resistance gene and overexpressed P450 enzymes that detoxify pyrethroids [20]. The *An. gambiae* s.l. Covè strain was also resistant to dieldrin while the *An. coluzzii* Akron strain was resistant to carbamates partly mediated by the Ace-1R mutation. However, we detected very low resistance ratios (<2 fold) to broflanilide in these strains compared to the susceptible Kisumu strain which suggests the absence of cross-resistance to broflanilide and the mechanisms of resistance to pyrethroids, dieldrin and carbamates they contain. Although broflanilide acts on the insect GABA receptor just like dieldrin, it’s site of action is different which may explain the lack of cross-resistance to broflanilide [10]. These findings together with the high susceptibility to broflanilide observed in high intensity pyrethroid resistant *An. gambiae* s.l. from 5 villages in Southern Benin would suggest that broflanilide could be very effective against a wide range of *An. gambiae* s.l. vector populations in the West African that exhibit similar insecticide resistance profiles. Further studies investigating the discriminating concentrations of broflanilide to other Anopheline vector species and cross resistance to broflanilide and other existing insecticide resistance mechanisms in malaria vectors are advisable.

## 5. Conclusion

In this study, we determined a discriminating concentration of 2.2 μg per bottle of broflanilide against *An. gambiae* s.l. vectors in bottle bioassays with the adjuvant Mero^®^ applied at 800 ppm. Using a lower dose of Mero^®^ (500 ppm) the discriminating concentration was 6 μg per bottle. Wild pyrethroid-resistant populations of *An. gambiae* s.l. from southern Benin were fully susceptible to the insecticide at both discriminating concentrations. There was no evidence of cross-resistance to broflanilide via mechanisms of resistance to pyrethroids, dieldrin or carbamates. Broflanilide thus showed potential to be highly effective against insecticide resistant *An. gambiae* s.l. malaria vector populations. VECTRON^™^ T500, a new wettable powder IRS formulation of broflanilide being assessed by WHO’s PQT/VCP team, has potential to improve vector control and enhance capacity to manage insecticide resistance if added to the list of pre-qualified IRS products. A large scale roll out of the insecticide for IRS might therefore be expected in the next few years. The discriminating concentrations determined in this study need to be confirmed at other trial facilities across Africa, but serve as a good starting point for monitoring susceptibility to broflanilide in Benin and making programmatic decisions around its deployment for improved vector control and insecticide resistance management.

## Declarations

### Consent for publication

Not applicable

### Availability of data and material

The datasets used and/or analysed during the current study are available from the corresponding author on reasonable request.

### Authors’ contributions

RG supervised the laboratory bioassays, analysed the data and prepared the graphs and tables. AO performed the dose-response assays, DT performed the insecticide susceptibility tests with insectary strains while AF performed the bottle bioassays with wild vector mosquitoes. GS contributed to manuscript revision. CN designed the study, acquired funding, supervised the project, and prepared the final manuscript.

## Acknowledgements

We thank Dr. Kunizo Mori of Mitsui Chemicals Agro, Inc for supplying the insecticide. We also thank the staff of CREC/LSHTM (Thomas Syme, Juniace Ahoga, Josias Fagbohoun, Estelle Vigninou, Abel Agbevo etc) for their assistance and Dr. Germain Gil Padonou, Director of CREC, for administrative support. We appreciate Drs Derric Nimmo, Sarah Rees and Ms. Janneke Snetselaar of IVCC for program support.

## Authors’ information

Not applicable

## List of abbreviations

IRS: Indoor Residual Spraying
WHO: World Health Organization
WHO PQT/VCP: World Health Organisation Vector Control Product Prequalification
(WHO PQT/VCP)GABA: γ -aminobutyric acid
WP: Wettable Powder
CREC: Centre de Recherche Entomologique de Cotonou
IVCC: Innovative Vector Control Consortium
LSHTM: London School of Hygiene & Tropical Medicine
Kdr: Knockdown resistance
Ace-1R: acetylcholinesterase insensitive mutation

